# deepNF: Deep network fusion for protein function prediction

**DOI:** 10.1101/223339

**Authors:** Vladimir Gligorijević, Meet Barot, Richard Bonneau

**Author notes:** Email address (Richard Bonneau).

## Abstract

The prevalence of high-throughput experimental methods has resulted in an abundance of large-scale molecular and functional interaction networks. The connectivity of these networks provide a rich source of information for inferring functional annotations for genes and proteins. An important challenge has been to develop methods for combining these heterogeneous networks to extract useful protein feature representations for function prediction. Most of the existing approaches for network integration use shallow models that cannot capture complex and highly-nonlinear network structures. Thus, we propose *deepNF*, a network fusion method based on *Multimodal Deep Autoencoders* to extract high-level features of proteins from multiple heterogeneous interaction networks. We apply this method to combine STRING networks to construct a common low-dimensional representation containing high-level protein features. We use separate layers for different network types in the early stages of the multimodal autoencoder, later connecting all the layers into a single bottleneck layer from which we extract features to predict protein function. We compare the cross-validation and temporal holdout predictive performance of our method with state-of-the-art methods, including the recently proposed method *Mashup*. Our results show that our method outperforms previous methods for both human and yeast STRING networks. We also show substantial improvement in the performance of our method in predicting GO terms of varying type and specificity.

**Availability:** *deepNF* is freely available at: https://github.com/VGligorijevic/deepNF

## 1. Introduction

Methods for automated protein function prediction allow us to maximize the utility of functional annotations derived from costly and time-consuming protein function characterization and large-scale genomics experiments. The accuracy of these methods has been improved with the advent of high-throughput experimental methods that have enabled construction of different types of genome-scale molecular and functional interaction networks, including protein-protein interaction networks, genetic interaction networks, gene co-expression networks and metabolic networks [2]. Extracting biological information from the wiring patterns (topology) of these networks is essential in understanding the functioning of the cell and its building blocks-proteins. A key insight behind this approach is that the function is often shared between proteins that physically interact [27], have similar topological roles in the interaction networks [18], or are part of the same complex or pathway [6].

Systematic benchmarking efforts, such as the Critical Assessment of Functional Annotation (CAFA)[25] and MouseFunc [23], has shown that the current state-of-the-art methods for protein function prediction use machine learning techniques to train classifiers on a multitude of network-, sequence- and structure-based data sources to make predictions [25, 23]. Due to the complementary nature of these different data sources, such techniques have been shown to be more accurate than those that use a single data source [16, 8, 31]. However, the heterogeneous nature of biological networks, as well as their different levels of sparsity and noise, make development of such techniques challenging. Here we focus on integrating only network-based features in order to limit the scope of the work, better isolate general results aimed at biological networks, and to better compare to important recent works on biological network integration.

Most previous approaches for network integration either use *probabilistic methods*, like Bayesian inference [11, 17] or *kernel-based methods* [34, 21] to fuse different protein-protein network types, derived from different proteomic and genomic data sources, into a single network. The resulting network, along with the set of proteins’ function labels, are fed into a kernel- or network-based classifier to derive functional associations of annotated proteins and generate hypotheses about unannotated proteins. For example, GeneMANIA [22, 20] is a widely used method that first integrates kernels of different network types into a single kernel by solving constrained linear regression problem; then, it applies Gaussian label propagation on the resulting kernel to make label predictions. However, as pointed out by [7], these methods suffer from the information loss incurred when combining all the network types into a single network. Moreover, they are not robust to noisy links and dense structures often present in certain network types (e.g., large blocks in gene co-expression networks or large hubs in protein-protein interaction networks); in this case, the resulting network is obscured by links from the noisy network types and can significantly impair the classification performance. To overcome these problems, some approaches train individual classifiers on these networks and then use *ensemble learning methods* to combine their predictions [35, 26, 32]. Lastly, such methods do not typically take into account correlations between different data sources, and often suffer from learning time and memory constraints. Previous work has also benefited from considering the hierarchical structure of Gene Ontology (GO) [10, 3], using statistical principles to choose negative examples (i.e., proteins without a given function) [33] or modeling the incomplete set of protein function annotations as a matrix completion or recommendation system problem [12].

## 2. Related work

A recent study proposed *Mashup* [7], a network integration framework, to address the challenge of fusing noisy and incomplete interaction networks. *Mashup* takes as input a collection of protein-protein association networks and applies a matrix factorization-based technique to construct compact low-dimensional vector representation of proteins that best explains their wiring patterns across all networks. These vectors are then fed into a Support Vector Machine (SVM) classifier to predict functional labels of proteins. The key step in *Mashup* is the feature learning step that constructs informative features that have been shown to be useful in multiple scenarios including highly accurate protein function and protein-protein interaction prediction.

There are several challenges to learning a useful low-dimensional network representation (also known as *network embedding*) while preserving the network structure. In particular, most protein-protein association networks are characterized by diverse connectivity patterns. Specifically, proteins with the same or similar functional annotations in these networks often exhibit a complex mixture of relationships, based both on homophily (close proximity to each other in the network) and structural similarity (similar local wiring patterns, regardless of the position in the network).

Thus, it is a challenging task to learn a lowdimensional embedding of proteins that preserves non-linear network structure while remaining predictive of protein function. Even more challenging is the construction of such a compact low-dimensional embedding of proteins that is consistent across different protein functional and molecular interaction modalities (i.e., across different types of protein-protein association networks).

The majority of previous network embedding methods use shallow and linear techniques that cannot capture complex and highly non-linear network structure. These include methods such as *node2vec* [13] and *DeepWalk* [24] that are mainly used on social networks. On the other hand, *deep learning* is a promising technique to deal with such problems, and has been shown to work well for problems such as speech recognition, natural language processing (NLP) and image classification, as well as for several biological problems [1]. Motivated by the recent success of deep learning techniques in learning powerful representations from complex data, a few recent studies propose using *deep neural networks (DNNs)* for computing network embeddings [13, 4, 30]. DNNs apply multiple layers of non-linear functions to map input data into a low-dimensional space, thereby capturing highly non-linear network structure in efficient low-dimensional features. A multi-layer architecture of DNN is a key to learning richer network representation. The advantage of using DNNs has been demonstrated in learning embeddings of large-scale social networks for performing different tasks, such as link prediction, network clustering and multi-label classification [30, 28, 4]. However, none of these methods can construct embeddings by handling different network modalities (i.e., types, views), i.e., these methods cannot be used for integrative analysis. Thus, we propose deep Network Fusion, *deepNF* (also pronounced *deep enough*), an integrative framework for learning compact low-dimensional feature representation of proteins that 1) captures complex topological patterns across multiple protein-protein association networks, and that 2) is used to derive functional labels of proteins. To explicitly address the diversity of protein-protein interaction network types, we use separate layers for handling each network type in the early parts of the deep autoencoder, later connecting all the layers into a single bottleneck layer from which we extract features to predict protein function for different species. Similar to *Mashup*, in the last phase, *deepNF* trains an SVM on the resulting features to predict each protein function label.

*deepNF* is based on a Multimodal Deep Autoencoder (MDA) to integrate different heterogeneous networks of protein interactions into a compact, lowdimensional feature representation common to all networks. An autoencoder is a special type of neural network that is composed of two parts: 1) an encoding part, in which the input data is transformed into low-dimensional features, and 2) a decoding part, in which those features are mapped back to the input data [29]. Our method, *deepNF*, has the following conceptual advances: 1) it **preserves the non-linear network structure** by applying multiple layers of non-linear functions, composing the DNN architecture of *deepNF*, thereby learning a richer network representation; 2) it **handles noisy links** present in the networks, as autoencoders have also been shown to be effective denoising systems capable of constructing useful representations from corrupted data [29]; and 3) it **is efficient and scalable** as it uses the MDA to learn low-dimensional protein features from all networks in a fully unsupervised way and independently of the function prediction task. This allows for the use of the entire data set in the training of the MDA, resulting in high-quality features. Our method enables semisupervised approaches to function prediction. Here we demonstrate such a semi-supervised approach to function prediction task by training an SVM for each function on these features. Additionally, the reduced dimension of the extracted features makes the training of the SVMs computationally efficient.

We apply this method on human and yeast STRING networks to construct a compact lowdimensional representation containing high-level protein features. For each species, we perform 5-fold cross validation, as well as temporal holdout validation, in which we train our method on GO annotations from 2015 and test it on those from 2017. We report the performance of our method for different DNN architectures. We contrast the performance of our method with the state-of-the-art network integration methods, *Mashup* and *GeneMANIA*. We report a significant improvement of *deepNF* over these methods on both yeast and human protein function annotations. We also report a significant improvement in performance when training *deepNF* on all STRING networks together than when training it on each individual STRING network, demonstrating the success of our integrative strategy.

To the best of our knowledge, this is the first method that uses a deep multimodal technique to integrate diverse biological networks. We demonstrate that deep learning methods offer the great advantage of being able to capture non-linear information contained in large-scale biological networks, and that using such techniques could lead to improved network representations. Features learned by using these methods not only lead to substantial improvements in protein function prediction accuracy but also our temporal holdout results indicate that our method can also be used for prioritizing novel experimental target proteins for a given function.

## 3. Approach

In this section we introduce our framework for predicting protein functions from multiple networks, *deepNF*, including the preprocessing step, adopted from [4], and the cornerstone of our method, the Multimodal Deep Autoencoder (MDA). In the preprocessing step the structural information of each network is converted into a high-dimensional vector representation that is used as input to the MDA. We also provide implementation details and a description of our testing schemes: cross-validation and temporal holdout validation.

## 4. Methods

We consider a set of *N* = 6 undirected weighted STRING networks whose connectivity patterns are represented by a set of symmetric adjacency matrices {**A**^(1)^, **A**^(2)^,…, **A**^(*N*)^}. Each matrix, 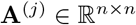, is constructed over the same set of proteins n. *deepNF* learns low-dimensional latent feature representation of n proteins, 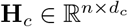 (where *d_c_* ≪ *n*), shared across all networks. In order to do so, the method follows three steps (see Fig. 1): (i) it converts structure of each network into a high-quality vector representation by first applying the *Random Walk with Restarts (RWR)* method and then constructing a *Positive Pointwise Mutual Information (PPMI)* matrix capturing structural information of the network; (ii) it fuses PPMI matrices of networks by using the MDA, and from the middle layer extracts a low-dimensional feature representation of proteins; (iii) it predicts protein function by training an SVM classifier on the low-dimensional features computed in the previous step. An outline of the procedure is provided in Algorithm 1. We provide details of each step below.

**Figure 1:**
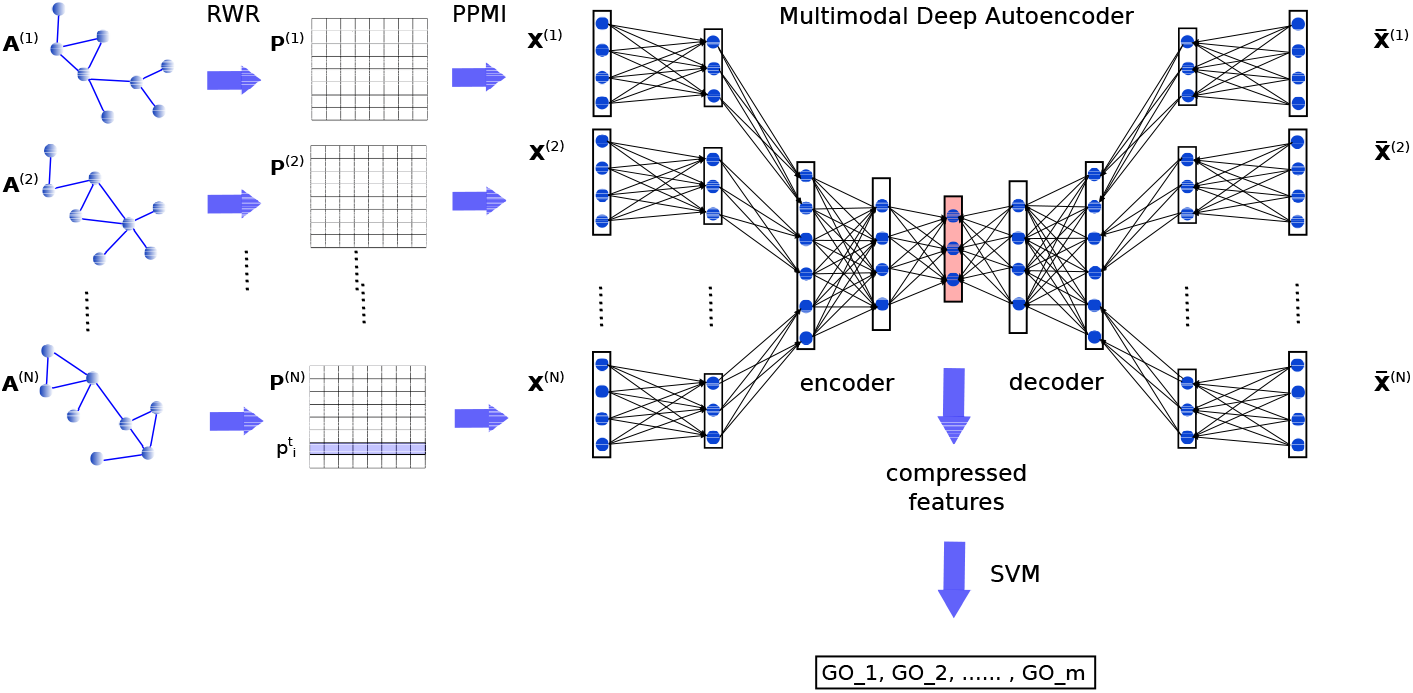
Method overview. In the first step networks are converted into vectors after the Random Walk with Restarts method (left). After this preprocessing step, the networks are combined via our MDA (right). Low-dimensional features are then extracted from the middle layer of the MDA and used to train a final classifier (bottom).

### 4.1. Random walk-based network representation

To capture network structural information and to convert it to high-dimensional protein vector representation suitable for input to the MDA, we adopt the approach of [4] and further modify it for multiple networks. For each network *j* ∈ {1,…*N*} we construct high-quality vector representations of proteins, 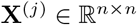, preserving potentially complex, non-linear relations among the network nodes. To do so, we first use the RWR model to capture network structural information and to characterize the topological context of each protein. We chose the RWR method for converting network structure into initial node vector representations over the previously proposed sampling-based procedure in *node2vec* [13] and *DeepWalk* [24], because these methods are computationally more intense and require additional hyperparameter fitting. The RWR approach can be formulated as the following recurrence relation:

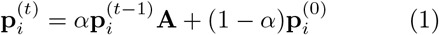

where 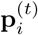 is a row vector of protein *i*, whose *k*-th entry indicates the probability of reaching the *k*-th protein after *t* steps, 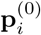 is the initial 1-hot vector, *α* is the probability of restart controlling the relative influence of local and global topological information of a network represented by adjacency matrix A. By expanding the iterations in the recurrence relation in equation 1, the probabilities of reaching to all proteins in a network after *T* steps can be written as:

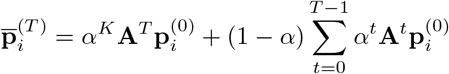

It has been shown that for constructing a good node representation, random walks of greater lengths should be given less weight in order to capture relevant local connections and reduce the introduction of noise coming from distant nodes of the network. Thus, the final representation of the *i*-th protein can be constructed in the following way: 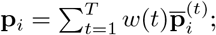 where *w*(*t*) is a monotonically decreasing *weight function* [13, 4, 19]. Here, we adopt the weighting strategy proposed by [4], which results in the following mathematical form used for computing node representations:

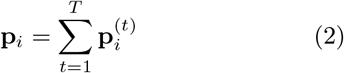

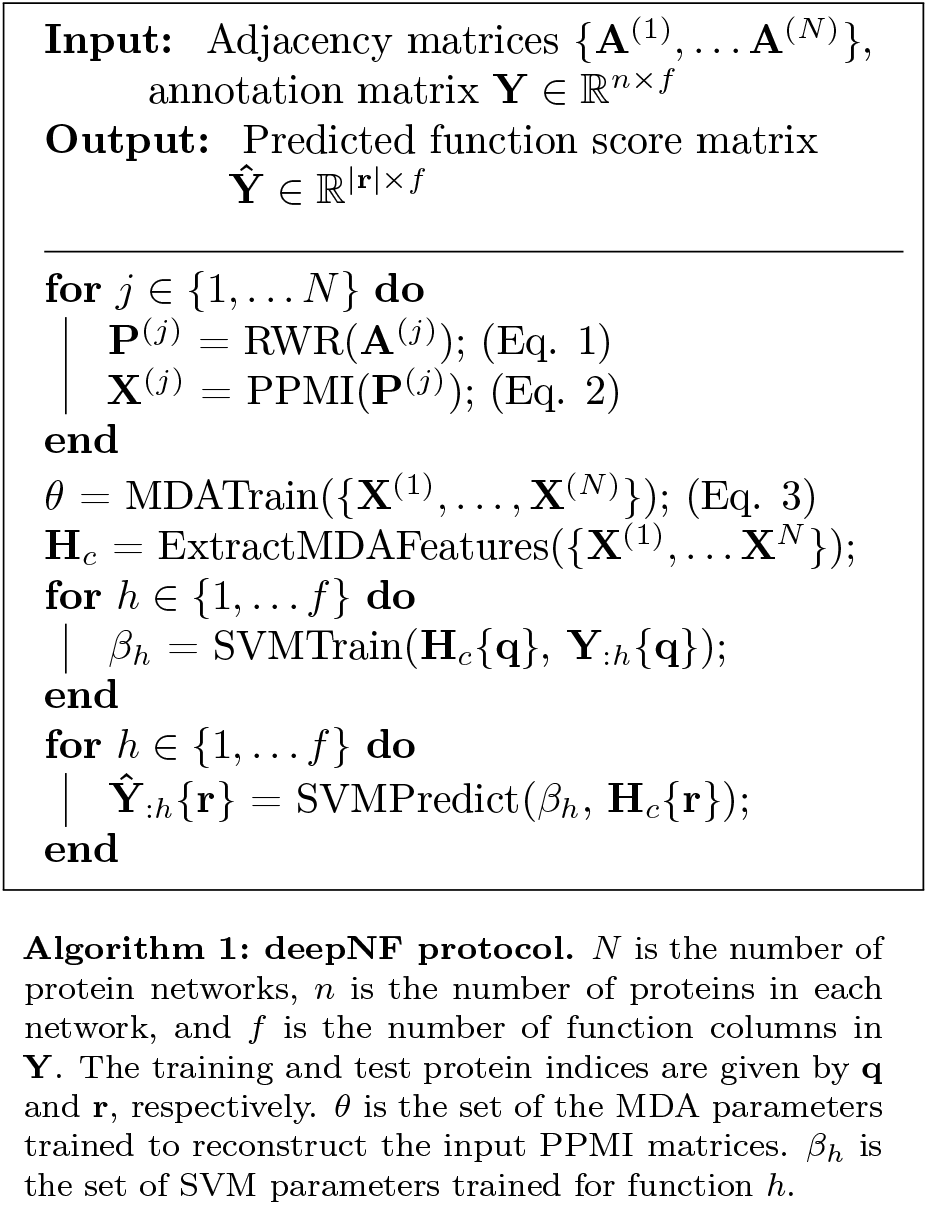

Repeating this process for every node *i* ∈ {1,…*n*} in the network *j*, results in a representation matrix 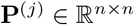, characterizing the probability of co-occurrence of network nodes. Next, from the probabilistic co-occurrence matrix, **P**^(*j*)^, of network, *j*, we construct a vector representation of proteins by computing PPMI matrix defined as:

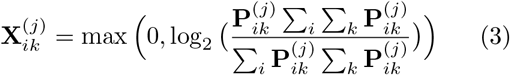

It is important to note that this process occurs as a first step and thus the RWR representation is mitigating the sparsity of some individual network types prior to the deeper integration described in next steps below.

### 4.2. Integrating networks with MDA

Although the above approach is fast, it results in protein features that still represent individual networks in the high-dimensional space. As such, these features cannot be readily used for protein function prediction. Here, we describe MDA, a novel method for integrating multiple networks represented by PPMI matrices, reducing their dimension and creating protein features, extracted from all networks, that are more suitable for training a classifier and predicting protein functions.

The MDA constructs a low-dimensional feature representation of n proteins, that best approximates all networks, *j* ∈ {1,…,*N*}, by projecting their PPMI matrices, **X**^(*j*)^, using multiple non-linear activation functions, into a common feature space, 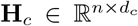 (i.e., a common bottleneck layer in DNN architecture of the MDA, see Fig. 1). Following the standard definition of autoencoders [29], we formulate the encoding and decoding part of the MDA as follows:

- *Encoding:* in the first hidden layer of the MDA, we first compute low-dimensional non-linear embedding, 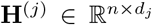, for each network *j* ∈ {1,…, *N*}:

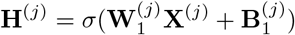

then, we compute a common feature representation by applying multiple non-linear functions (i.e., by stacking a series of hidden layers in the MDA) on the feature representation obtained by concatenating features from all networks obtained in the previous step (i.e., the previous layer):

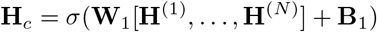
- *Decoding:* we compute reconstructed PPMI matrices, 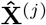, for each network, *j* ∈ {1,…, *N*}, by mapping the common representation, 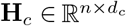, first to the lower dimensional representations of the individual networks, and then back to the original space, by also applying multiple non-linear functions:

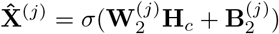

where, 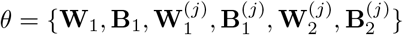 is the set of all parameters in both the encoding and decoding parts of our model to be learned in the training process, 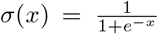 is the *sigmoid* activation function and [*] denotes concatenated matrices.

The aim of the MDA method is to find optimal 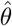 that minimizes the reconstruction loss, *L*(*θ*), between each original and reconstructed PPMI matrix:

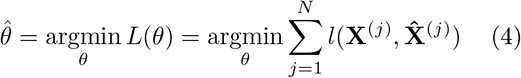

where *l*(*) is the sample-wise *binary cross-entropy function*.

A key step of our approach is the second step in the encoding part of the MDA that constructs a common feature representation by first denoising each individual network, by constructing their corresponding low-dimensional feature representations, and then projecting them into a common feature space.

The loss function (equation 4) can be optimized by standard back-propagation algorithm. We use minibatch stochastic gradient descent with momentum for training the MDA. We also explore the performance of the MDA with different batch sizes, learning rates, and different architectures (i.e., number and sizes of hidden layers). Values of all hyperparameters are provided in Section 3 of the Supplementary Material. After the training of the MDA is done, we extract the low-dimensional features, 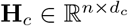, from its bottleneck layer.

### 4.3. Predicting function from multiple networks

We model the problem of protein function prediction as a multi-label classification problem. We use the compressed features, **H**_*c*_, computed in the previous step, to train an SVM classifier to predict probability scores for each protein. We use the SVM implementation provided in the LIBSVM package [5]. To measure the performance of the SVM on the compressed features, we adopt two evaluation strategies: (a) *5-fold cross validation* and (b) *temporal holdout validation*.

In the *5-fold cross validation*, we split all annotated proteins into a training set, comprising 80% of annotated proteins, and a test set, comprising the remaining 20% of annotated proteins. We train the SVM on the training set and predict the function of the test proteins. We use the the standard radial basis kernel (RBF) for the SVM and perform a nested 5-fold cross validation within the training set to select the optimal hyperparameters of the SVM (i.e., *γ* in the RBF kernel and the weight regularization parameter, *C*) via grid search. All performance results are averaged over 10 different CV trials.

In the *temporal holdout validation*, we use the protein GO annotations from 2015 and 2017 to form training, validation and test sets. We form the training set from proteins whose annotations did not change from 2015 to 2017. We form the test set from the proteins that did not have any annotations in 2015 but gained annotations in 2017. We use the same setup for the SVM classifier as in 5-fold cross validation. To fit the hyperparameters of the SVM, we created a validation set comprising proteins who had annotations in 2015 but also gained new annotations in 2017. We choose the optimal hyperparameters based on the SVM performance on the validation proteins, and report the final results on the test proteins. The performance results are averaged over 1,000 bootstraps of the test set.

We compare the performance of our method with 2 state-of-the art methods, *Mashup* and *GeneMANIA*. For each method, we apply the validation strategies described above. We use the following metrics to evaluate the prediction performance: (i) *Accuracy (ACC)*, that measures the percentage of test proteins that were correctly predicted (i.e. a protein is correctly predicted if the set of its predicted functions exactly match the set of its known functions); (ii) *Micro-averaged F1 score (F1)* is computed in the same way as in [7]; (iii) *Micro-averaged area under the precision-recall curve (m-AUPR)* is computed by first vectorizing the protein-function matrices of predicted scores and known binary annotations, and then computing the area under the precision-recall curve by using from these two vectors; *Macro-averaged area under the precision-recall curve (M-AUPR)* is computed by first computing the AUPR for each function separately, and then averaging these values across all functions. Here, we do not consider receiver operating characteristic (ROC) curves as protein labels are highly skewed, and AUPR is less biased in that case and thus better choice [9].

### 4.4. Data preprocessing

To make the *cross-validation* performance comparison of *deepNF* with *Mashup* fair, we use the exact same dataset (i.e., the six STRING networks and functional annotations) used in the Mashup paper [7]. The basic network measures and properties of the STRING networks are provided in Table S1 in the Supplementary Material. Also, we report the results of the methods on both yeast and human STRING networks. The functional annotations for yeast are taken from Munich Information Center for Protein Sequences (MIPS) (again, this is done to make a perfect comparison to validations performed in [7]) and they are organized into three functional categories: Level 1 (consisting of 17 most general functional categories), Level 2 (consisting of 74 functional categories) and Level 3 (consisting of 153 most specific functional categories). All functional annotations for human are taken from GO. Similar to yeast, they were arranged into three functional categories, i.e., categories containing GO terms annotating 11-30 (covering 153 molecular function, MF, 262 biological process, BP, and 82 cellular component, CC GO terms), 31-100 (covering 72 MF, 100 BP and 48 CC GO terms) and 101-300 (covering 18 MF, 28 BP and 20 CC GO terms) proteins respectively.

To compile function annotation data for *temporal holdout* validation, we use the same strategy proposed in the CAFA challenge [25, 15]. We obtain protein annotation data for 2015 (release 145) and 2017 (release 167) year from UniProt-GOA [14] database. For each ontology (i.e., MF, BP and CC) and each model organism (i.e., yeast and human), we create our training, validation and test sets of proteins as described above, specifically: the training set is formed of proteins whose annotations did not change from 2015 to 2017, the test set comprises proteins who did not have any annotations in 2015 and gain at least 10 new annotations in 2017, and the validation set comprises proteins that had annotations in 2015 but also gained new annotations in 2017. We consider only GO terms that were gained by test proteins from 2015 to 2017 and that have between 10 and 300 training proteins in 2015. The number of training, validation, and test proteins, as well as the number of new functions for MF, BP and CC, for yeast and human, are summarized in Table S2 in the Supplementary Material.

## 5. Results

Here, we use cross-validation and temporal holdout to evaluate *deepNF* and compare its performance to *GeneMANIA* and *Mashup*. In all of our experiments we set *α* = 0.98 in the RWR step of our method as this leads to the best generalized results across all function label types and architectures tested for both organisms. Other choices of α (e.g., α = 1.0) have been shown to result in lower quality of extracted features (see Sec. 4.1) [4]. In the training of the MDA, we explore different layer configurations (also known as architectures) and regularization parameters. Values of all the hyperparameters and the details of the MDA training strategy are provided in Sec. 3 in the Supplementary Material.

### 5.1. Cross-validation performance

To evaluate the quality of the low-dimensional features, extracted from the bottleneck layer of the MDA (Fig. 1), we run the same five-fold crossvalidation procedure as in the *Mashup* paper [7]. We train the MDA for different layer configurations for Yeast and Human STRING networks. The performance of our method in yeast and human, for different architectures, is provided in Figs. S1 and S3 in the Supplementary Material, respectively. We find that the features obtained from the 5-layer architecture (2 encoding, 1 feature layer and 2 decoding layers) of the MDA, trained on the Yeast STRING networks, leads to the best performance in terms of the m-AUPR across all three levels of MIPS ontology. Performance of the same model on different annotation levels of the MIPS hierarchy, in comparison to *GeneMANIA* and *Mashup*, is summarized in Fig. 2. We observe that *deepNF* significantly outperforms (rank-sum p-value < 0.01) both *Mashup* and *GeneMANIA* in terms of m-AUPR at different levels of MIPS hierarchy. Consistent improvement of *deepNF* is also achieved in terms of accuracy (i.e., the percentage of test proteins with all the predicted functions exactly matching the corresponding known functions). Namely, *deepNF* accurately assigns known functions to 31.3% of proteins, as opposed to 23.6% for *GeneMANIA* and 25.5% for *Mashup* in Level 1 of MIPS annotations. Note that this is more rigorous measure than the accuracy measure used in the *Mashup* paper by [7] that only considers top predicted functions for each protein. Although we observe clear improvements in terms of m-AUPR across all levels of MIPS annotations, however, in terms of M-AUPR in Levels 2 and 3 of MIPS annotations, *deepNF* performs comparably to *Mashup*. We have a highly unbalanced multi-label problem and M-AUPR aggregates the contributions of all GO terms with equal weights; thus, M-AUPR is not as suitable a measure for unbalanced labels as m-AUPR. Thus, we reason that the consistently higher values of m-AUPR across different levels of yeast MIPS ontology that we observe here indicates that our method can handle the unbalanced labels much better than the other methods.

**Figure 2:**
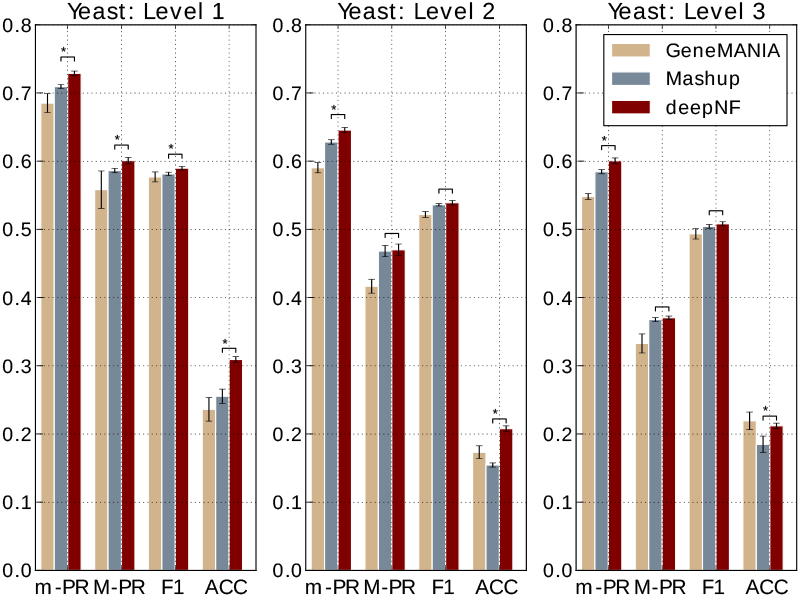
Cross-validation performance of our method in integrating yeast networks. Performance of our method, with the MDA architecture [6 × *N*, 6 × 2000, 600], in 5-fold cross validation in comparison to function prediction performance of the state-of-the-art integration method, *Mashup*, and *GeneMANIA*. Performance is measured by the area under the precision-recall curve, summarized over all GO terms both under the micro-averaging (m-PR) and macroaveraging (M-PR) schemes; F1 score and accuracy (ACC). Performance of the methods is shown separately for MIPS yeast annotations for Level 1 (left), Level 2 (middle) and Level 3 (right). The error bars are computed based on 10 trials. Asterisks indicate where the performance of *deepNF* is *significantly* higher than the performance of *Mashup* (rank-sum p-value < 0.01).

The cross-validation performance of the 7-layer MDA (3 econding, 1 feature layer and 3 decoding) applied on Human STRING networks in comparison to *Mashup* and *GeneMANIA* is shown in Fig. 3. Our method significantly outperforms the other two methods, in terms of all four measures, for the MF-GO terms belonging to the most general (i.e., annotating between 101-300 proteins) and the most specific (i.e., annotating between 11-30 proteins) categories. The performance of our method for MF-GO terms with between 31-100 proteins annotated in the training set) is comparable to *Mashup*, except in terms of F1 measure for which our method achieves significantly better performance. Similar results are also observed for both BP and CC ontologies (shown in Fig. S5 in the Supplementary Material). The observed improvement in accuracy of our method in comparison to *Mashup* can be partially attributed to the high quality of protein features extracted from the complex topology of STRING networks in the hierarchical manner. Unlike *Mashup*, which utilizes a shallow matrix factorization-based technique to construct compact protein feature representation, *deepNF* utilizes a hierarchical way of feature construction by incorporating intermediate layers in the MDA architecture (see Fig. 1); features constructed in this way capture fine-grained topological patterns in the large-scale STRING networks. The de-noising property of the multimodal autoencoder, underlying our method, leads to better detection of relevant features from individual networks and ultimately to a better final integrated feature representation. To further demonstrate the usefulness of such an approach in feature construction, we also apply our method on individual STRING networks (i.e., without integration). Namely, we train a deep autoencoder on each STRING network, separately, and further assess the quality of the extracted lowdimensional features of each individual network in predicting protein functions. The integrative performance and the performance on individual networks of our method in comparison to *Mashup* is shown in Fig. 4, for both yeast and human STRING networks.

**Figure 3:**
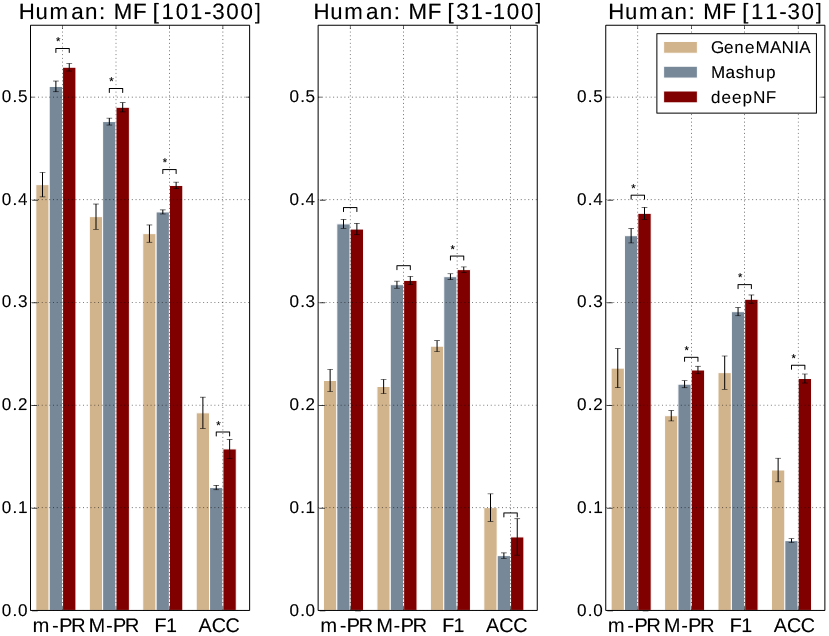
Cross-validation performance of our method in integrating human networks. Performance of our method, with the MDA architecture [6 × N, 6 × 2500, 9000,1200], in 5-fold cross validation in comparison to function prediction performance of the state-of-the-art integration method, *Mashup*, and *GeneMANIA*. Performance is measured by the area under the precision-recall curve, summarized over all GO terms both under the micro-averaging (m-AUPR) and macro-averaging (M-AUPR) schemes; F1 score and accuracy (ACC). Performance of the methods is shown separately for all three ontologies of GO, i.e., MF, BP and CC, where each ontology is further divided into three levels annotating 101-300, 31-100 and 11-30 proteins respectively.

### 5.2. Temporal holdout performance

Unlike the cross-validation procedure, that randomly divides protein set into folds used for training and testing the model, the temporal holdout procedure divides proteins into training and test sets based on their annotations at two different widely-separated time points where older annotations are used for training and newer ones are used for testing the model. The temporal holdout approach ensures a more “realistic” scenario of function prediction. The study of individual MDA architectures shows that the 5-layer architecture of the MDA in yeast and 7-layer architecture of the MDA in human yields the best performance in terms of M-AUPR across different GO ontologies (see Figs. S2 and S4 in the Supplementary Material). The temporal holdout validation performance of our method with these architectures is shown in Figs. 5 and 6, for yeast and human data, respectively. The performance of both methods on molecular function terms is higher than for biological process terms, which is in line with previous studies [25, 15].

**Figure 4:**
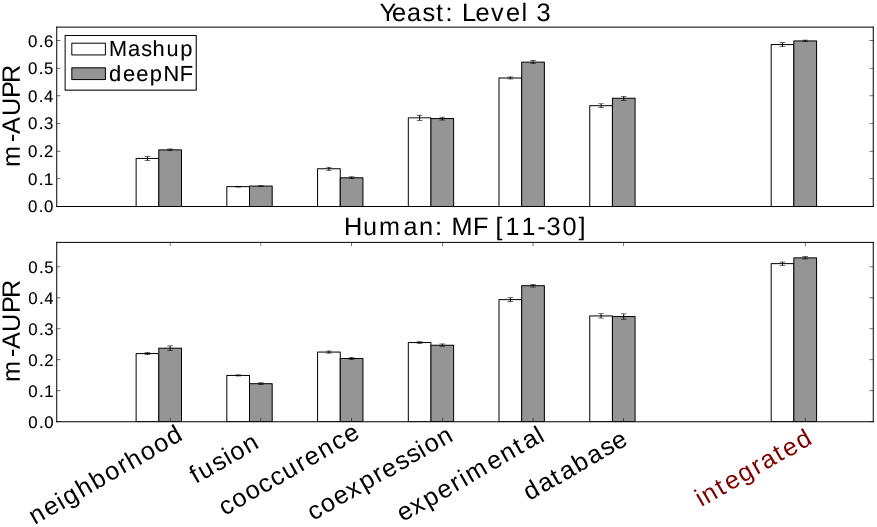
Integrating multiple networks outperforms individual networks in protein function prediction. We compare the cross-validation performance of our method applied on individual STRING networks, measured by m-AUPR, with the performance of *Mashup*. The upper panel shows the performance results on the most specific MIPS terms (Level 3) for each individual STRING network of yeast, whereas, the bottom panel shows the performance results on the most specific MF-GO terms for each individual STRING network of human. The low-dimensional features of these networks are extracted from the bottleneck layer of autoencoders trained on each individual network. We use architecture [*N*, 2000, 600] for yeast STRING networks, and architecture [*N*, 2500, 9000,1200] for human STRING networks. In addition to individual network performance, we also show the integrative performance of both methods.

**Figure 5:**
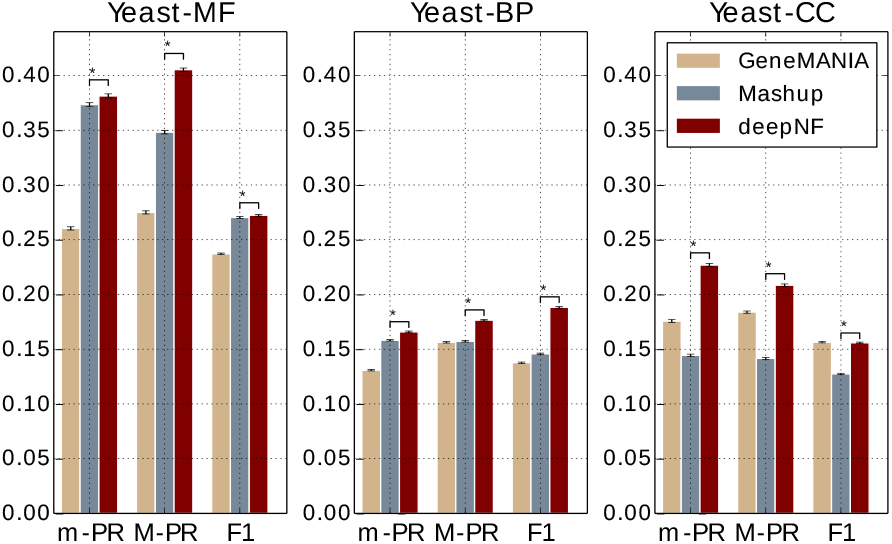
Temporal holdout validation performance of our method in integrating Yeast STRING networks. Performance of our method with the MDA architecture [6 × *N*, 6 × 2000, 600], in temporal holdout validation in comparison to function prediction performance of *GeneMANIA* and *Mashup*. Performance is measured by the area under the precision-recall curve (AUPR) both under micro-averaging (m-AUPR) and macro-averaging (M-AUPR) and F1 score. The results are averaged over 1,000 boost-raps of the test set; asterisks indicate where the performance of *deepNF* is *significantly* better than the performance of *Mashup* (rank-sum p-value < 0.01).

**Figure 6:**
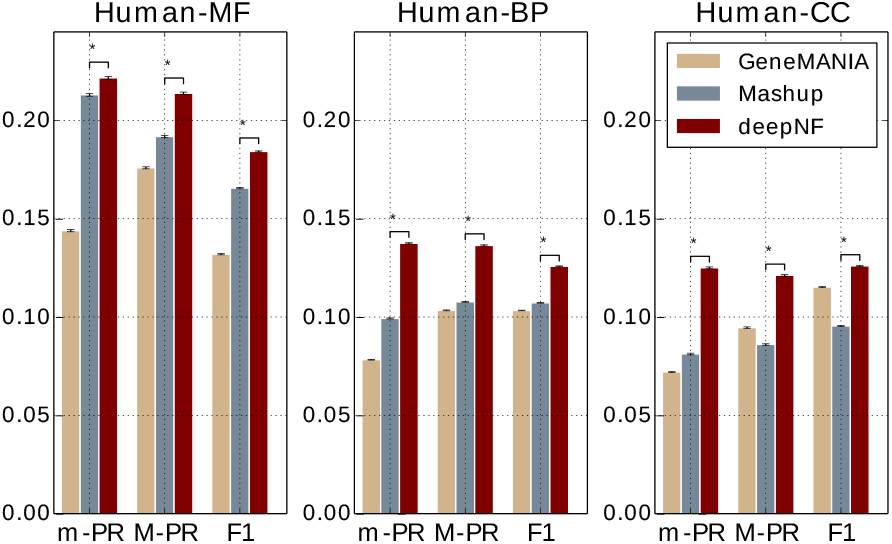
Temporal holdout validaton performance of our method in integrating Human STRING networks in comparison to GeneMANIA and Mashup. Performance of our method, with the MDA architecture [6 × *N*, 6 × 2500, 9000, 1200], in temporal holdout validation in comparison to function prediction performance of *GeneMANIA* and *Mashup*. Performance is measured by the area under the precision-recall curve (AUPR) both under micro-averaging (m-AUPR) and macro-averaging (M-AUPR) and F1 score. The results are averaged over 1,000 boost-raps of the test set; asterisks indicate where the performance of *deepNF* is *significantly* better than the performance of *Mashup* (rank-sum p-value < 0.01).

We observe that *deepNF* substantially outperforms both *Mashup* and *GeneMANIA* in temporal holdout validation. We observe clear improvement in both yeast and human data across all three types of ontologies. Interestingly, unlike in cross-validation, where *Mashup* significantly outperforms *GeneMANIA*, in temporal holdout validation, especially for the cellular component (CC) ontology for both yeast and human data, *GeneMANIA* achieves higher performance results than *Mashup*. This could be due to the very high density of CC annotations (i.e., there are on average 2.42 CC-GO terms per protein in the training set out of total 11 CC-GO terms, as opposed to 2.54 MF-GO terms out of total 20 MF-GO terms and 3.77 BP-GO terms per protein out of total 43 BP-GO terms, in the temporal holdout set in yeast), that can be handled better by label propagation framework, such as *GeneMANIA*, than by *Mashup*. However, the high density of annotations per protein is more suitable for a deep learning technique, such as our method, that achieves higher performance, across all metrics, than both *GeneMANIA* and *Mashup*.

We further explored the performance of our method on specific individual GO terms used in the temporal holdout study. The AUPR values of the 43 BP-GO terms computed from the features of Yeast STRING networks are shown in Fig. 7. From the figure, we can observe that *deepNF* achieves higher AUPR performance than *Mashup* for the majority of BP-GO terms. We observe similar results also for MF- and CC-GO terms (see Fig. S7 in the Supplementary Material), and also for Human STRING networks in all three ontologies (see Fig. S8 in the Supplementary Material). Specifically, we observe that majority of MF-GO terms associated with “binding” perform better with deep architecture (i.e. *deepNF*) than with a shallow one (i.e., *Mashup*), whereas, the quite opposite situation is observed with MF-GO term associated with “transporter activity”. Surprisingly, this is in contrast with one of the findings reported in [25], where the authors observe that most function prediction methods consistently perform worse on the “binding” related terms than on the “transporter activity” related terms. This indicates that our method provides complementary results in comparison to the methods presented in the paper. Furthermore, by looking into CC-GO terms, we observe that majority of CC-GO terms associated with “membrane” perform better with our method than with *Mashup*, whereas the situation is opposite for CC-GO terms associated with “vesicle”.

**Figure 7:**
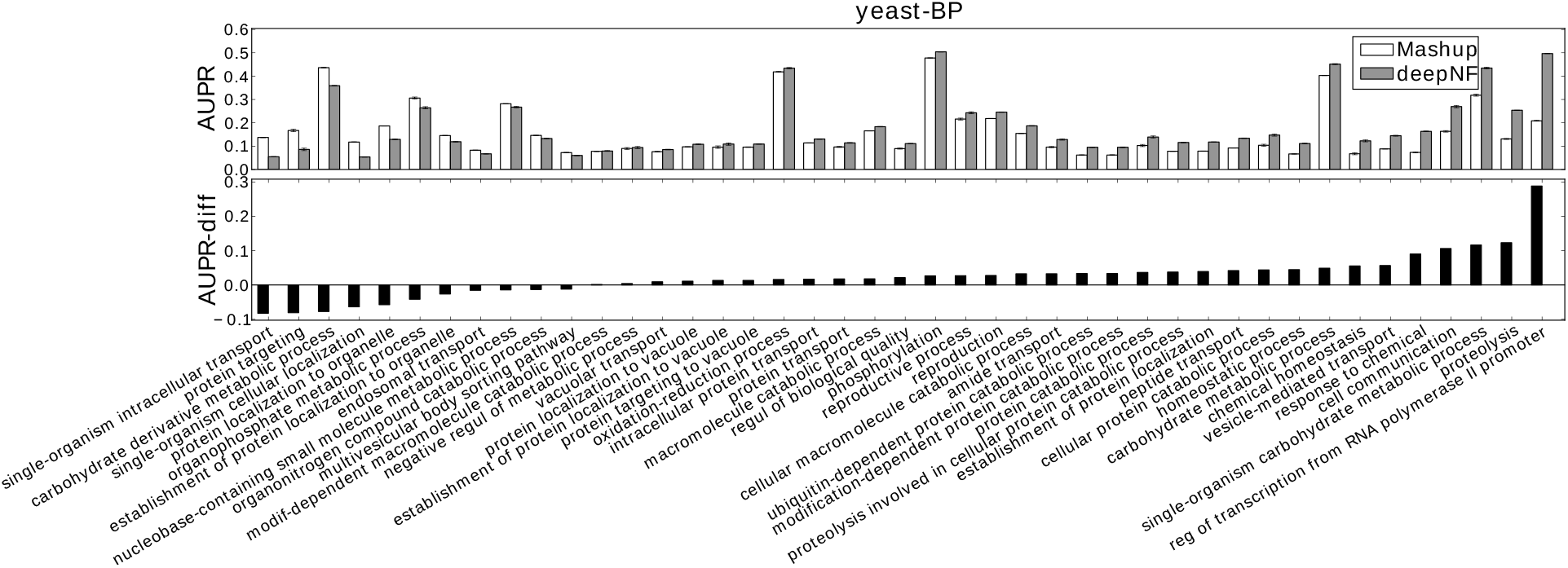
Temporal holdout validation shows significant difference in the performance of our method vs. *Mashup* in predicting individual yeast BP-GO terms. For each BP-GO term we show the *Mashup*’s and *deepNF*’s performance (5-layer MDA), measured by the AUPR. The lower panel shows the difference in the performance of these two methods. The names of the GO terms are shown on the x-axis.

## 6. Conclusion

Recent wide application of high-throughput experimental techniques has provided complex highdimensional complementary protein association data; the wide availability of this data has in turn driven a need for protein function prediction methods that can take advantage of this heterogeneous data. We present here, for the first time, a deep learning-based network fusion method, *deepNF*, for constructing a compact low-dimensional protein feature representation from a multitude of different networks types. These features allow us to use out-of-the-box machine learning classifiers such as SVMs to accurately annotate proteins with functional labels.

*deepNF* extracts features that are highly predictive of protein function, which is attributed to the fact that the method relies on a deep learning technique that can more accurately capture relevant protein features from the complex, non-linear interaction networks. Unlike *Mashup* (an innovative previous method that combines protein networks to generate features for function prediction) cannot extract features that have this quality.

We present an extensive performance analysis comparing our method with competing protein function prediction methods. In addition to crossvalidation, the analysis includes a temporal holdout validation evaluation similar to the measures in Critical Assessment of Functional Annotation (CAFA). Double-blind field-wide validation efforts (like CAFA) have demonstrated the utility of such temporal holdouts and established them as the most accepted way of performance comparison for protein function prediction methods. We show that *deepNF* outperforms previous methods (*Mashup* and *GeneMANIA)* in both human and yeast organisms, in multiple levels of specificity of gene ontology and MIPS terms.

Given that the features generated by *deepNF* are task-independent, they can be used for other applications besides protein function prediction. Additionally, our method is not limited to only network integration: in future work, we hope to explore integrating non-network information such as protein sequences and structures into our representations in order to make more accurate predictions of protein function.

## Acknowledgements

The authors would like to thank Da Chen Emily Koo for enlightening discussions and help with construction of the temporal hold-out validation sets. We thank Ian Fisk, Nicholas Carriero and Dylan Simon of the Simons Foundation for discussion and help with high performance computing.

## Funding

The authors acknowledge the support of the Simons Foundation, the NIH, the NSF and NYU for supporting this research, particularly NSF: MCB-1158273, IOS-1339362, MCB-1412232, MCB-1355462, IOS-0922738, MCB-0929338, and NIH: 2R01GM032877-25A1.

